# Assessing the robustness of human ncRNA notations

**DOI:** 10.1101/2024.12.08.627405

**Authors:** Nadia K. Prasetyo, Paul P. Gardner

## Abstract

The HUGO Gene Nomenclature Committee (HGNC) is the only worldwide authority that assigns standardised nomenclature to human genes (1). All studies related to the human genome and genes worldwide should adhere to HGNC-approved gene names and symbols, emphasizing the importance of precise classification and naming. Recent studies have revealed the functional and clinical relevance of RNU2-2P, which is linked to neurodevelopmental disorders and cancer (2–4), underscoring the need to reassess the classification of pseudogenes and functional non-coding RNA genes. In this study, we explore the conservation and expression of genes from 15 small ncRNA families, including U1, U2, U4, U5, U6, U4ATAC, U6ATAC, U11, U12, Vault tRNA (VTRNA), Y RNA, tRNA, 7SL, U7, and 7SK, to identify non-coding RNA-derived pseudogenes that are under strong negative selection in the human genome. Our findings highlight three highly conserved and expressed pseudogenes—RNU2-2P, RNU1-27P, and RNU1-28P—that are likely misclassified, as existing evidence suggests they may play a role in disease research. This warrants a reevaluation of their status as pseudogenes. Additionally, we identified RNU5F-1, a functional copy of RNU5, which is lowly conserved and expressed, yet its classification as a functional gene raises questions about its potential role. Furthermore, other pseudogenes and functional ncRNAs that could also be misclassified were identified, suggesting the necessity for further experimental and clinical examination.

## Introduction

Pseudogenes are defined as non-functional sequences derived from their functional gene counterparts. They have high sequence similarity to one or more gene paralogues but differ to them at crucial points, rendering them non-functional, rapidly degraded, untranscribed or untranslated (for protein-coding genes) (5). However, there is evidence that some pseudogenes are transcribed and are not transcriptional junk (6), and may have some biochemical functions (7–10). Hence, the discrimination between functional genes and pseudogenes is a hard problem. Even more so for non-coding RNA (ncRNA) pseudogenes, the identification of ncRNA pseudogenes is further a challenge. Some excellent results have been obtained by evaluating structure-based covariance models and sequence-based hidden Markov models in tRNAs (11). The difference of which gives an indication of how strongly selection is acting to maintain structure. However, this approach breaks down for unstructured ncRNAs (e.g. C/D box snoRNAs) and likely those RNAs with dynamic structures (e.g. riboswitches, spliceosomal RNAs). Separately, computational analysis of transcriptomics and epigenetics has also been used to identify transcribed pseudogenes from RNA-Seq data (12). A combination of these techniques may offer a method to distinguish pseudogenes from genes by analysing structure, expression patterns, and conservation (13).

Next generation sequencing and whole genome studies have highlighted the importance of ncRNA variation in the human genome, enabling the discovery of associations between ncRNAs and diseases. Recent studies have been focused on the clinical significance of ncRNAs, including in cancer and genetic disorders (14,15). Latest research highlights the critical role of highly conserved spliceosomal snRNAs, revealing de novo variation in RNU4-2, RNU2-2P, and RNU5B-1 are associated with neurodevelopmental delays and related disorders (2,3,16,17). A further 50 snRNA were investigated and further disease associations discovered (18). An interesting detail emerged with the variation found in RNU2-2P, annotated as a U2 snRNA pseudogene (as of 19/11/2024, updated late November 2024) in major human genome databases such as the human genome nomenclature (HGNC) database (19). The evolutionary conservation tracks from UCSC show high levels of conservation of RNU2-2P, higher even than RNU2-1. Indicating strong levels of negative selection are maintaining this sequence across the millions of years since it appeared in the genome (Figure 1). This raises the question: How reliable are the pseudogene notations provided by HGNC, and could there be other “functional” pseudogenes linked to diseases that remain unidentified due to their classification as pseudogenes, also are some “functional” ncRNAs more likely to be pseudogenes?

**Figure 1:**
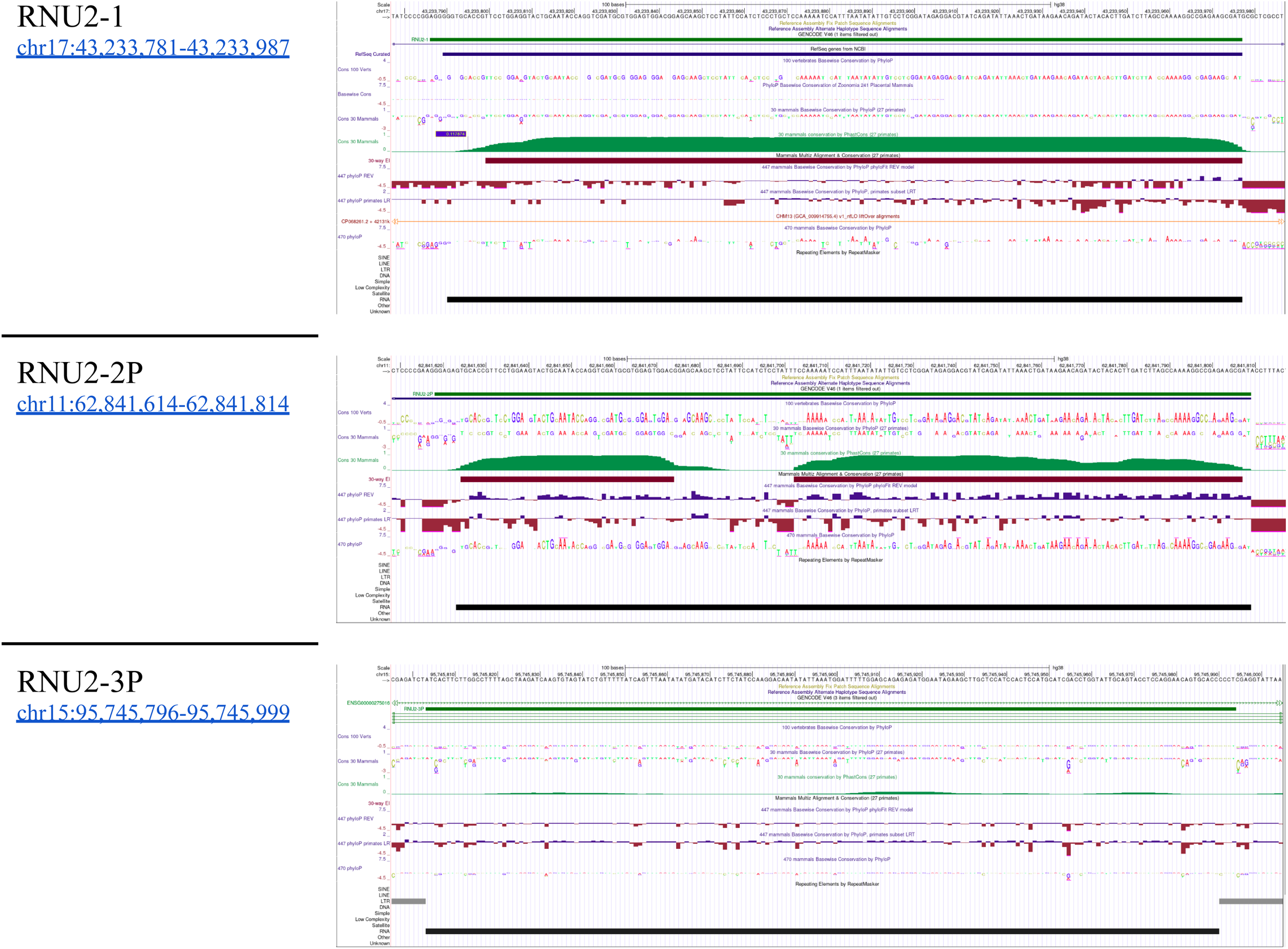
Evolutionary conservation of the U2 spliceosomal RNA (RNU2-1), the U2 pseudogene of interest (RNU2-2P) and a control U2 pseudogene (RNU2-3P).

In this study, we compare the classification of ncRNA genes (and ncRNA-derived pseudogenes) in the HGNC database with evolutionary conservation and transcription information. The aim is to identify pseudogenes and ncRNAs that are outliers in terms of conservation across primates, mammals and vertebrates, along with gene expression data of each gene. Those that are under strong negative selection in the human genome and highly expressed are likely to be functional, providing a means to reclassify potential mis-classifications. These may be important for further disease research and the status of outlier pseudogenes and ncRNAs should be reconsidered.

## Materials and Methods

### ncRNA Dataset

The non-coding RNA data used in this study was derived from approved gene symbols from the HGNC database ((20) last updated 2024-10-25). The gene symbols found for a specific RNA were inspected using Ensembl Release 113 (21) to extract chromosomal coordinates for each gene symbol. To verify the location and access alignments and annotations, the datasets for each ncRNA were mapped to the UCSC genome browser. Several gene symbols identified in HGNC lacked associated transcript IDs and were subsequently queried and mapped to the UCSC genome browser HGNC annotations.

### Assessment of conservation of pseudogenes

The evolutionary conservation status for each ncRNA and pseudogene was collected from phastCons30 (27 primates), phyloP100 (100 vertebrates), and phyloP447 (477 mammals) annotations in the UCSC Genome Browser as a bigwig files (<u>phastCons30way</u>, <u>phyloP100way</u>, and <u>pyloP447wayBW</u>). PhastCons30 contains single nucleotide conservation scores based on individual alignment columns and its flanking columns (22); (23) whilst the phyloP measures conservation at individual nucleotides, ignoring the effects of their neighbours. Conservation scores were compiled and analysed in R version 4.4.1 to assess the concordance between ncRNA pseudogene annotations and conservation in comparison to functional ncRNA copies.

### Assessment of expression of genes and pseudogenes

Gene expression data were derived from the NIH Genotype-Tissue Expression (GTEx) project, available to download from the UCSC database (<u>GTEX RNA Seq Coverage</u>), and ENCODE project (24,25), available from the ENCODE portal (<u>ENCODE RNA-Get</u>). Expression was determined in 52 tissues and 2 cell lines, on 17,382 samples from 948 adults. The ENCODE expression included total, polyA plus, and polyA minus RNA-seq databases on all biosample classifications (77 human tissue, 52 human cell lines, 210 human primary-cell samples, and 36 In vitro differentiated cell samples) (<u>Matrix - ENCODE</u>). The maximum FPKM across conditions for each ncRNA and pseudogene was collected.

### Analysis of highly conserved and expressed RNA pseudogenes

Kolmogorov-Smirnov (KS) tests were conducted for each group of genes, as well as for pooled gene groups, to evaluate differences in conservation between functional ncRNAs and their pseudogenes. In addition to this, a principal component analysis (PCA) plot was generated to detect any outliers in the classification of pseudogenes and functional ncRNAs. The analysis revealed that the two numerical features most strongly distinguishing these gene categories are PhyloP100 vertebrate conservation scores and ENCODE gene expression tracks. These features demonstrated the highest KS D statistic (maximum difference) values and the most statistically significant P values, highlighting their importance in differentiating between functional ncRNAs and pseudogenes.

### Robust z-scores for PhyloP100 and ENCODE analysis

To standardise the PhyloP100 and ENCODE data to comparable scales, we calculated robust z-scores. These scores facilitate the consistent evaluation of differences between functional ncRNAs and pseudogene controls for each feature. We selected robust z-scores that rely on the median and median absolute deviation (MAD), rather than the mean and standard deviation based standard z-score which is sensitive to outliers and skewed distributions. However, when the MAD is at or near zero, this calculation becomes problematic. In such cases, we substitute the MAD with the “mean absolute deviation” (MnAD). The following formulas were applied (26):

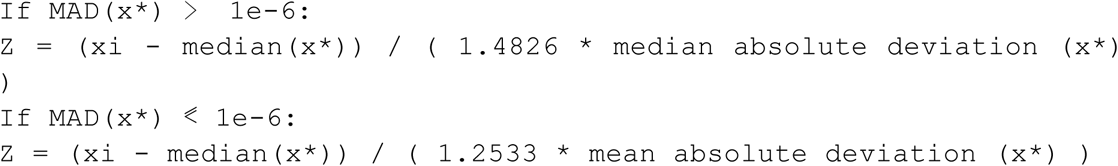

Here, x* represents the pseudogene control datasets of each gene group, ensuring that all datasets are scaled consistently. This approach improves visualisation by aligning axes comparably and supports analyses of different data types.

### Random Forest Algorithm

To estimate the likelihood of a gene’s functionality, we developed a random forest algorithm that evaluates two factors: the PhyloP100 conservation median and the ENCODE maximum expression for each gene, categorizing them into functional ncRNAs and pseudogenes. The model was trained using data from 476 functional genes and 2,618 pseudogenes, and tested on a smaller set of 52 functional genes and 292 pseudogenes. The classification model demonstrated good performance on the test dataset, achieving a macro-averaged F1-score of 0.92, and a ROC AUC score of 0.99, indicating excellent precision and recall. The model was then used on 5 ambiguous pseudogenes, one ambiguous functional gene, and one functional gene control, generating a functional probability for each ncRNA.

### Data Availability

Noncoding RNA and nomenclature data for the selected pseudogenes of the Human Genome: Dec. 2013 (GRCh38/hg38) are available from the HGNC website (https://www.genenames.org/), and UCSC genome browser (https://genome.ucsc.edu/cgi-bin/hgGateway). Gene symbols not found in Ensembl were searched using HGNC annotations available from the UCSC database as a bigbed file (<u>HGNC (7.8 M)</u>) last modified 2023-02-20 17:11. Conservation data is available to download as a bigwig file in <u>phastCons30way(6.6 G)</u>: last updated 2017-11-06 08:52; <u>phyloP100way</u> <u>(9.2G)</u>: last updated 2015-05-08 15:37; <u>pyloP447wayBW (9.3G)</u>: last updated 2023-08-30 12:18. Ensembl Release 113 data was accessed through two primary methods: BioMart service and library to connect and access the “hsapiens_gene_ensembl” dataset for human genes and the Ensembl REST API to access the location data including the chromosome, start, and end positions of each transcript. Both of these are available in the Ensembl web page for public use, with specified request per second limits (21). GTEx gene expression data are available as 54 individual tissue expression data, each of which are separate bigwig files in <u>GTEX RNA Seq Coverage (19.522G)</u>, last updated 2019-12-16 09:20. ENCODE gene expression data are available on the ENCODE portal v134.0 (<u>ENCODE RNA-Get</u>).

### Code Availability

The analysis of the snRNA data was done using Python: version 3.10.12, and R version 4.4.1. The code used in the analysis is available from https://github.com/NadiaPrasetyo/ncRNA-pseudogenes. R packages used include: ggplot2 version 3.5.1, BiocManager version 1.30.25, readr version 2.1.5, dplyr version 1.1.4, ggrepel version 0.9.6, stringr version 1.5.1. Python packages used include: biomart v0.9.2, mysql-connector-python v9.1.0, numpy v2.1.3, pip v22.0.2, pyBigWig v0.3.23, requests v2.32.3.

## Results

### Highly Conserved ncRNA Pseudogenes

To determine the highly conserved ncRNA pseudogenes, three sets of conservation data were collected: PhyloP100 (Vertebrate conservation), PhyloP447(Mammalian conservation), and PhastCons30 (Primates conservation). Kolmogorov-Smirnov (KS) test was performed on pooled small ncRNA divided into functional gene and pseudogene groups. The small ncRNA included genes and pseudogenes from U1, U2, U4, U5, U6, U4ATAC, U6ATAC, U11, U12, Vault tRNA (VTRNA), Y RNA, tRNA, 7SL, U7, and 7SK families (Table 1).

**Table 1:**
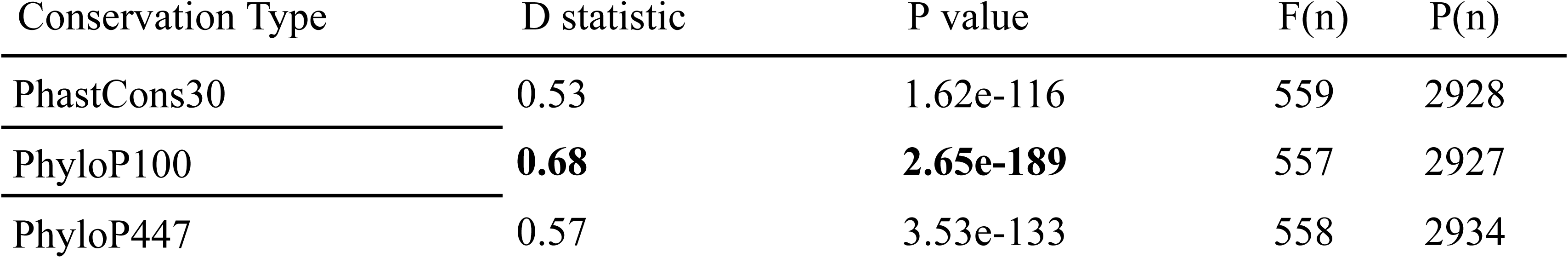
Kolmogorov-Smirnov test summary of the three conservation types. The D statistic is the maximum difference between the empirical distribution function of functional and pseudogene. The P value outlines the probability of obtaining the results observed (P value < 0.05 is significant). F(n) is the number of functional ncRNA gene samples, and P(n) is the number of ncRNA pseudogene samples.

| Conservation Type | D statistic | P value | F(n) | P(n) |
| --- | --- | --- | --- | --- |
| PhastCons30 | 0.53 | 1.62e-116 | 559 | 2928 |
| PhyloP100 | **0.68** | **2.65e-189** | 557 | 2927 |
| PhyloP447 | 0.57 | 3.53e-133 | 558 | 2934 |

PhyloP100 conservation showed the highest D statistic and lowest P value, indicating high and significant difference between functional and pseudogene ncRNAs within the included ncRNA families.

To identify highly conserved pseudogene outliers, a robust z-score normalisation method was used to scale the PhyloP100 conservation data and jitter-plots produced for gene families, grouped by functional ncRNA and pseudogene (Figure 2). Pseudogenes with z-scores higher than 2 were considered significant, indicating conservation more than 2 median absolute deviation (MAD). Pseudogenes with z-scores > 3 signify likely extreme outliers including RN7SKP70, RNU6-1334P, RNU6-1194P, TRL-TAA5-1, RNU2-2P, RNU5B-5P, and RNU6-1059P. Some functional ncRNAs also had low z-scores (z-score < 0) including the U5 “functional” gene RNU5F-1. Comparison of the pooled functional and pseudogenes (Figure 3) shows distinct difference in median z-score of functional (med = 2.90) and pseudogenes (med = 0). There are many highly conserved pseudogene outliers, along with functional ncRNA genes with low conservation values (Supplementary Table 1).

**Figure 2:**
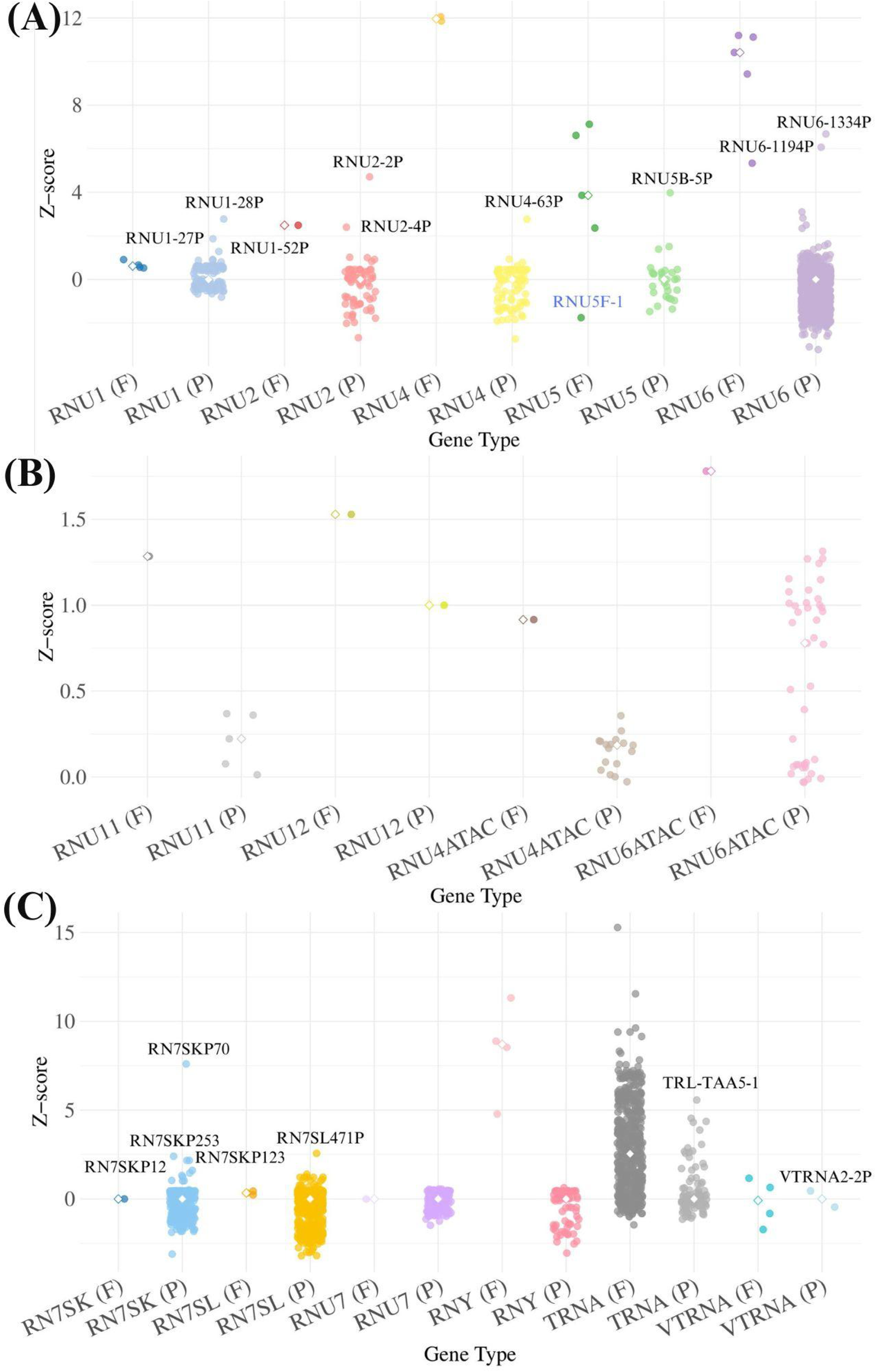
Z-score normalised vertebrate conservation (PhyloP100) of **A)** Major spliceosomal RNA, **B)** Minor spliceosomal RNA, and **C)** Other small ncRNA. Outliers of each gene family pseudogenes are labelled in black; outliers of functional ncRNAs within its gene family are labelled in blue. The complete list of outliers excluded from this plot is in Supplementary Table 1.

**Figure 3:**
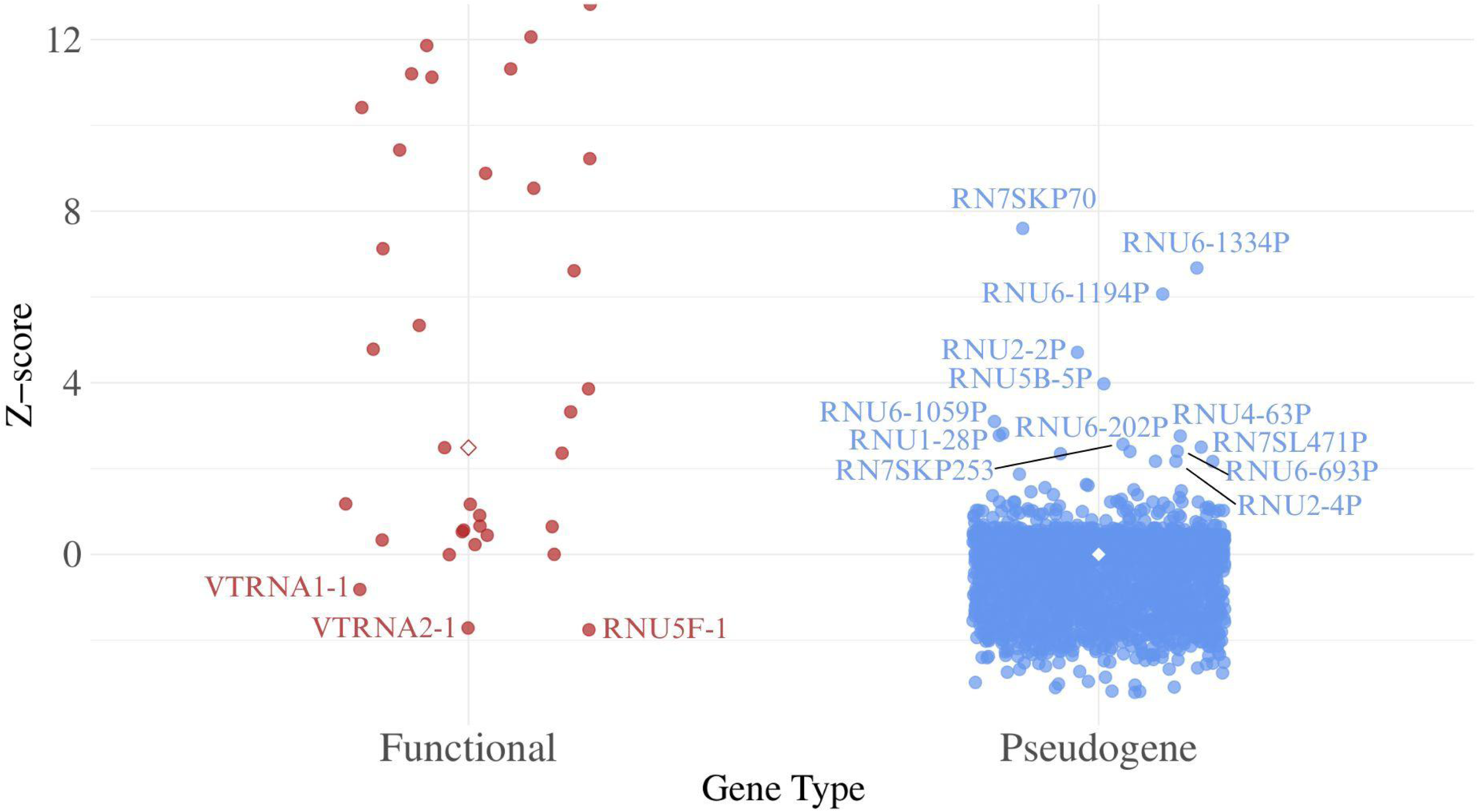
Z-score normalised vertebrate conservation (PhyloP100) of pooled small ncRNA families excluding tRNA, grouped into functional ncRNA and pseudogene. Pseudogenes are coloured blue, and highly conserved pseudogenes are labelled in blue. Functional ncRNA genes are coloured red, and low conserved functional ncRNAs are labelled. The complete list of PhyloP100 outliers are listed in Supplementary Table 1.

### Highly Expressed ncRNA Pseudogenes

To analyse the gene expression levels of ncRNA functional ncRNAs and pseudogenes, ENCODE total, poly-A plus, and poly-A minus RNA-seq data from human samples were retrieved via the ENCODE RNA-Get portal (25). The maximum expression level across samples for each ncRNA and pseudogene was extracted and compared using a KS test to assess differences between functional ncRNA and pseudogenes. This analysis yielded a KS D statistic of 0.68 and a P-value of 7.78e-23, based on data from 534 functional ncRNA genes and 2959 pseudogene samples across the ncRNA families. Robust z-scores were then computed to quantify the deviation of each ncRNA gene’s maximum expression from the pseudogene median. These values were visualised on a logarithmic scale (Figure 4). Notably, some pseudogenes exhibited z-scores above 3, reflecting abnormally high expression levels for some pseudogene. Furthermore, pseudogenes such as RNU1-27P, RNU1-28P, RNU2-2P, and various RNU6 and RNU7 pseudogenes, surpassed the expression levels of their corresponding functional ncRNAs.

**Figure 4:**
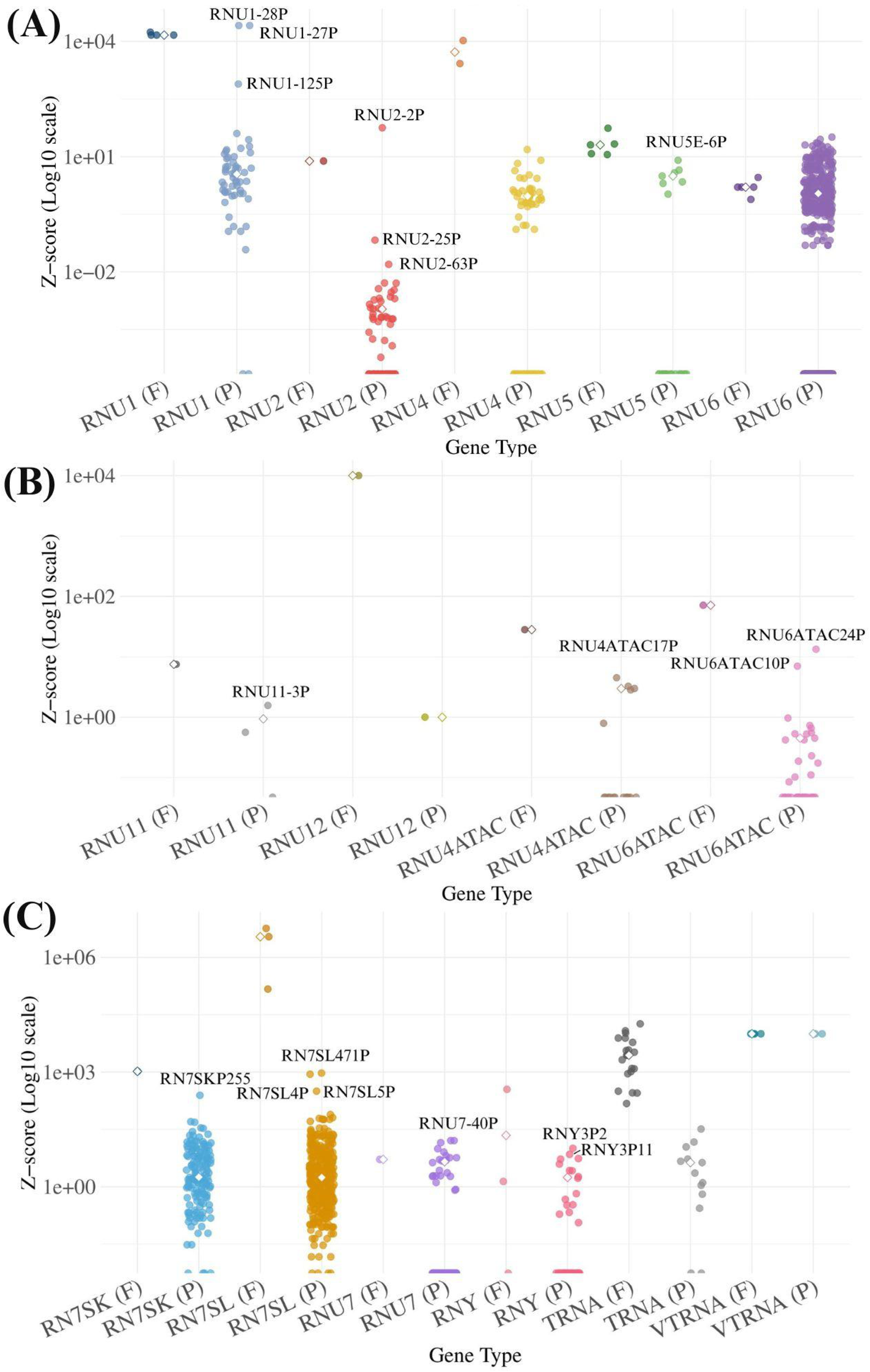
Z-score normalised gene expression (ENCODE RNA-seq) for **A)** Major spliceosomal RNA, **B)** Minor spliceosomal RNA, and **C)** other small ncRNA. Outlier pseudogenes within each gene family are labelled in black. Genes with a z-score of 0 are positioned at the bottom of the graph. Supplementary Table 2 provides the full list of outliers excluded from this plot.

The pooled functional ncRNAs and pseudogenes were compared to visualise differences in their distributions and identify outliers (Figure 5). A significant distinction was observed between the medians, with functional ncRNA genes having a median of 9.99 and pseudogenes a median of 0, highlighting a notable disparity in expression levels. Pseudogene outliers with exceptionally high expression levels border the pseudogene - functional gene boundary. Some functional ncRNAs exhibited low expression levels, comparable to those of pseudogenes.

**Figure 5:**
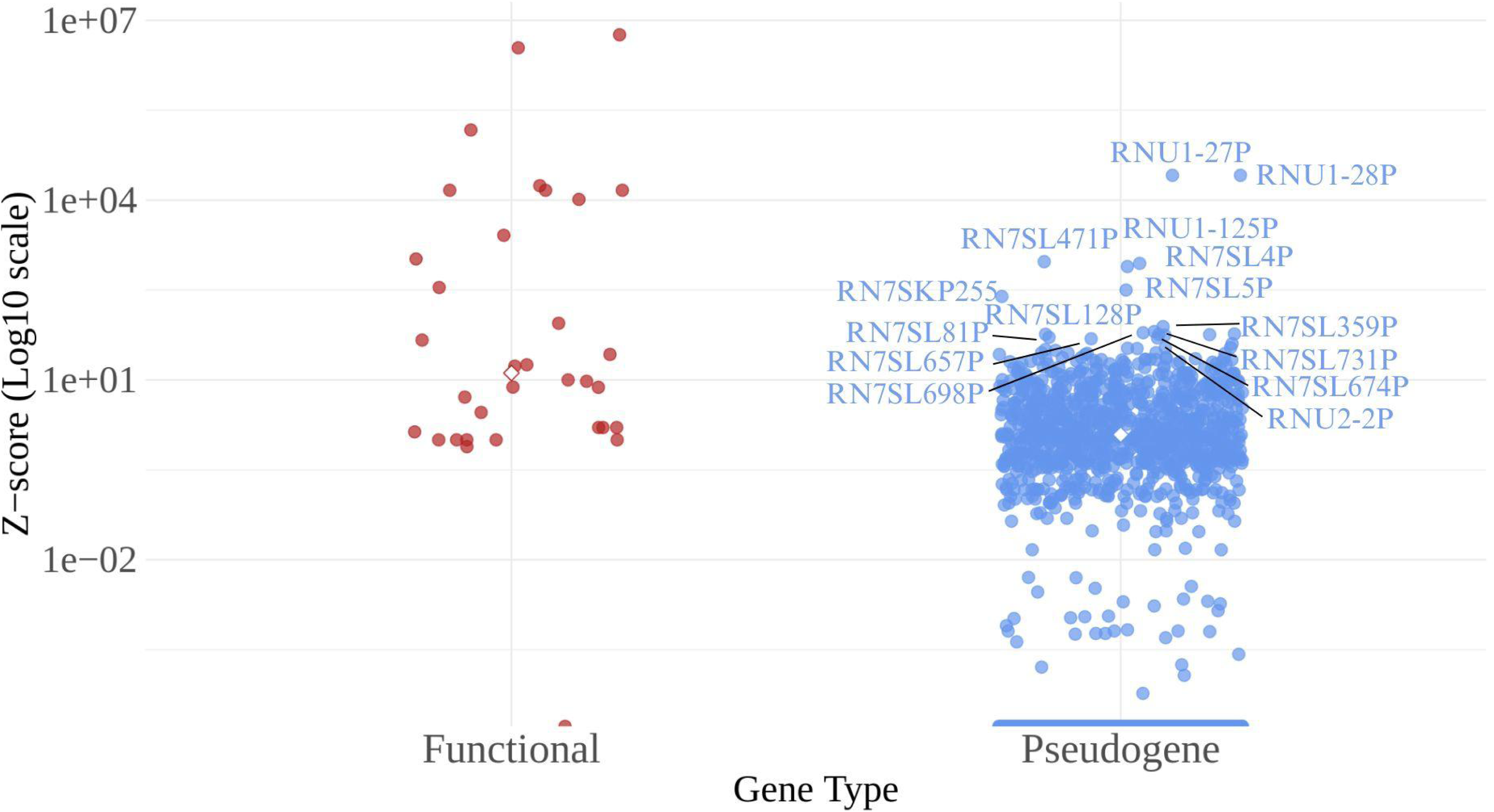
Z-score normalised gene expression (ENCODE RNA-seq) data for pooled small ncRNA families excluding tRNA, categorised into functional ncRNA and pseudogene. Pseudogenes are depicted in blue, with highly conserved ones specifically labelled. Functional ncRNA genes are shown in red, and those with low conservation are labelled accordingly. Genes with a z-score of 0 are located at the bottom of the graph. A comprehensive list of ENCODE outliers can be found in Supplementary Table 2.

### Functional Pseudgoenes and Non-functional Functional ncRNA

To identify annotation inconsistencies, we analysed the relationship between maximum expression levels and the median vertebrate conservation for all ncRNA genes (Figure 6A). The expression data (from ENCODE) were normalised to a standard distribution for clearer visualisation and comparison with vertebrate conservation. The scatter-plot reveals that gene functionality can be indicated by either high expression or high conservation, even if these traits are independent. Moderate levels of both further support functionality, with the combination of high conservation and high expression being the strongest indicator of functionality. Pseudogenes generally show low conservation with some expression (potentially noise) or high conservation with little to no expression, but outliers with moderate expression and conservation levels are associated with functional elements. Notable examples include RNU6-1189P, RNU6–28P, RNU2-2P, RNU1-27P, RNU5F-1 and RNU1-28 with abundant expression and moderate evolutionary conservation. To further test the likelihood of functionality, a random forest model was created and trained using the numerical features: maximum expression and vertebrate conservation (Figure 6B). The model was used to assess the functional probability of notable genes, providing the following predictions for ambiguous genes: RNU1-27P (1.00), RNU1-28P (1.00), and RNU2-2P (1.00), suggesting these genes are highly likely to be functional. The known functional control RNU5B-1 received a probability of 0.85, while RNU5F-1, RNU6-1189P, and RNU6-82P had probabilities of 0.00, indicating they are likely pseudogenes. These results highlight the anomalous behavior of RNU2-2P, RNU1-27P, RNU1-28P, and RNU5F-1, which may have been misclassified.

**Figure 6:**
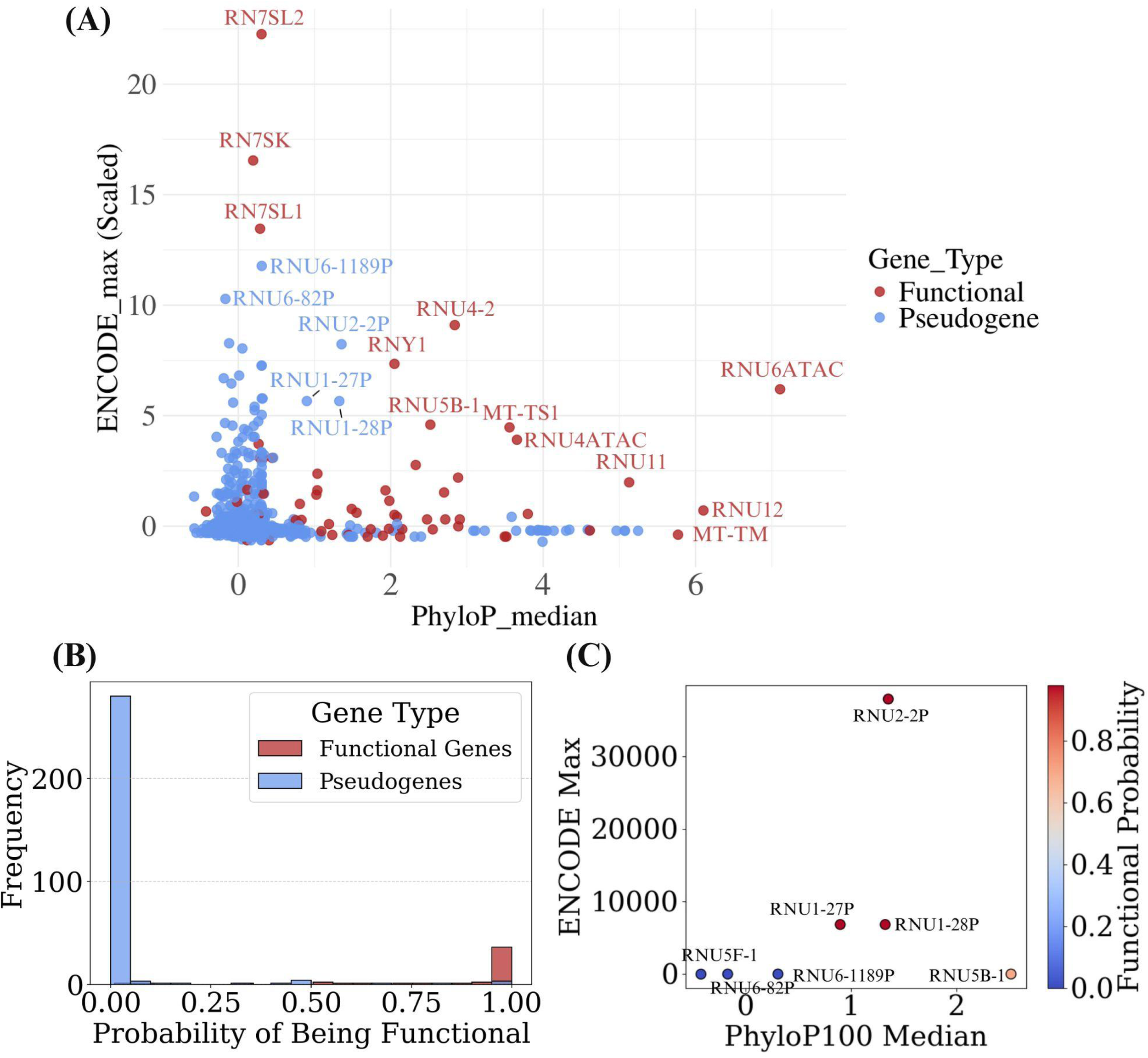
**A)** Scatter plot of maximum expression (ENCODE_max) and median vertebrate conservation (PhyloP_median) of all ncRNA genes. The Y-axis is scaled to a normal distribution of maximum expression levels to enhance visualisation. **B)** Distribution histogram of predicted functional probabilities for functional ncRNAs and pseudogenes in the test dataset. **C)** Scatter plot of predicted functional probability of RNU6-1189P, RNU6–28P, RNU2-2P, RNU1-27P, RNU1-28, and a functional control RNU5B-1. Functional gene points are coloured and labelled in red, whilst pseudogenes are coloured and labelled in blue.

## Discussion

Through the investigation of the conservation and expression data of the selected small non-coding RNA (ncRNA) families, the outlying pseudogenes with high conservation and expression have been identified. The pseudogene outliers have clear differences in expression and conservation levels from typical pseudogenes of each family (Figure 2, Figure 4).

The RNU1 family has 2 major outliers with high conservation and expression: RNU1-27P and RNU1-28P, both of which have significantly high z-scores for expression and high predicted functional probability (Figure 6A, 6C). U1 spliceosomal RNA is one of the small nuclear RNA (snRNA) in the major spliceosomal ribonuclease complex that is responsible to recognise and base pair to the 5’ of the splice site (27). Some recent findings show implications of recurrent U1 mutation in some multiple cancers (28,29) including mutations in RNU1-27P and RNU1-28P. This supports the probable functionality of RNU1-27P and RNU1-28P, with potential clinical variants significant in disease studies.

RNU2 has one specific pseudogene outlier, RNU2-2P that is highly expressed and conserved, even higher in both parameters than its functional copy RNU2-1 (Figure 1, 2A, 4A). U2 snRNA forms a ribonucleoprotein (U2 snRNP) that interacts with the 3’ region of the intron. RNU2 and RNU2-2P has gain popularity recently with many evidence pointing that RNU2-2P is likely functional, implicated in neurodevelopmental disorders and cancers such as Chronic lymphocytic leukaemia (CLL) and prostate adenocarcinoma (PRAD-UK) (2–4,30). Along with a 1.0 predicted functional probability (Figure 6C), it is very likely that RNU2-2P has been misclassified as a pseudogene.

RNU5 is another snRNA of the major spliceosome that forms a tri-snRNP assembly with the U4/U6 snRNP, creating the pre-catalytic B complex (27). The function of the U5 snRNP is dependent on the U5 snRNA loop, essential for the catalytic steps of aligning the 5’ and 3’ end of the exons (31). U5 has 5 functional copies: RNU5A-1, RNU5B-1, RNU5D-1, RNU5E-1, and RNU5F-1. While all five RNU5 copies contribute to splicing, they vary in their criticality and conservation. RNU5B-1, for example, has been implicated in various neurodevelopmental disorders, highlighting its clinical significance in maintaining proper splicing and cellular function (3,18). In contrast, RNU5F-1 shows much lower expression and conservation, more resembling a pseudogene (Figure 4A). This suggests RNU5F-1 may be inactive or only expressed in very specific conditions (10). Consistent with this, the random forest model assigns RNU5F-1 a low functional probability score of 0.0 (Figure 6C), indicating it likely has little role in most splicing and may have been misclassified as a functional gene.

RNU6, unlike the other major spliceosomal snRNAs, have numerous pseudogene copies (32). U6 has 5 functional copies, RNU6-1, 2, 7, 8, 9 and 1277 pseudogene copies annotated in the HGNC database. The results of conservation and expression analysis of all RNU6 copies show that while most of the pseudogenes are likely just pseudogenes, some might be functional. There are a large number of RNU6 pseudogenes with high conservation and expression listed in supplementary tables 1 and 2. Additionally, RNU6-1 has been found to be differently distributed in human carcinoma tissues, and an RNU6 pseudogene RNU6-505P differentially expressed in schizophrenic samples amongst others (33,34), signifying clinical diagnosis importance in the classifications of some of the RNU6 pseudogenes.

The minor spliceosome, formed from U11, U12, U5, U4atac, and U6atac splices around 0.35% of introns in the human genome. Although there are no highly conserved pseudogenes of any of the minor spliceosome snRNAs (Figure 2B), there are some that are highly expressed (Figure 4B), notably in RNU4ATAC and RNU6ATAC (Supplementary Table 2). RNU4ATAC has been found to be associated and important in many diseases including Microcephalic osteodysplastic primordial dwarfism type I (MOPD I), Taybi-Linder syndrome (TALS), Joubert syndrome, and many others (35–40). RNU4ATAC pseudogenes that are highly expressed including RNU4ATAC11P, RNU4ATAC17P, RNU4ATAC16P, RNU4ATAC18P may have potential disease significance and functional importance. Some RNU6ATAC pseudogenes are also found to be highly expressed (Figure 4B), with potential functions similar to U6atac mRNA regulation by rapid turnover acting as a real time transcriptional activity sensor (41). Although the clinical significance of RNU6ATAC has not been fully studied and understood, it is important to correct any misclassifications to provide the most accurate data for clinical and diagnosis research, preventing research exclusion of important variants.

This study expands the analysis of small ncRNA families by including additional ncRNA functional ncRNAs and pseudogenes. Most of these families have numerous copies, encompassing both functional ncRNAs and pseudogenes. Except for tRNAs, these small ncRNAs have only recently attracted research interest and understanding (42). As a result, the data quality and functional knowledge surrounding other small ncRNA families, such as 7SL, 7SK, Vault RNA, U7, and Y RNA, remain limited. Consequently, distinguishing the functional potential of these pseudogenes is likely inaccurate and largely speculative.

For instance, 7SL snRNA includes three HGNC approved functional ncRNAs and 684 identified pseudogenes. While many 7SL pseudogenes exhibit high expression levels (Supplementary Table 2), only one, RN7SL471P, shows significant conservation. However, given the role of 7SL in immunostimulation and signal recognition (42–44), the possibility that RN7SL471P has a functional role is a noteworthy finding.

7SK RNA exhibits a similar trend, with numerous pseudogenes showing high expression levels but only a few with significant conservation (Supplementary Table 1, 2), such as RN7SKP70 and RN7SKP253. Despite having just a single functional copy, 7SK RNA has been implicated in transcriptional regulation, cellular differentiation, and senescence (45–48). The presence of potentially functional 7SK RNA pseudogenes highlights the possibility of additional roles and importance of 7SK RNAs in the human body.

U7 snRNA was first identified as histone pre-mRNA 3’ processing factor by base pairing (49). Recent studies have shown that mutations in RNU7-1, the only functional copy of U7, are linked to Aicardi-Goutières syndrome (AGS) (50,51). Furthermore, the U7 snRNP complex, composed of U7 snRNA and associated proteins have been explored as a biotechnological tool in gene therapy targeting diseases caused by splicing defects (52,53)RNU7 pseudogenes with potential functionality, such as those exhibiting high expression levels (Supplementary Table 2), could play an important role in advancing gene therapy techniques and improving disease diagnosis.

Y RNAs are highly conserved in vertebrates and play crucial roles in initiating chromosomal DNA replication and regulating apoptosis (54–56). All four functional Y RNA copies have been linked to tumor development and cancer, with potential applications as cancer biomarkers and as molecular targets for anti-proliferative therapies (57). RNY pseudogenes with high expression levels (Supplementary Table 2) may possess previously unrecognized roles and functions, potentially contributing to cancer development.

Vault RNAs (VTRNA) and transfer RNAs (tRNA) present unique patterns of conservation and expression. VTRNAs have unknown function (42,58), while tRNAs have more functional copies than pseudogenes, unlike other small RNA families. These characteristics make it challenging to confidently identify functional pseudogenes within the VTRNA and tRNA families. The absence of highly conserved and highly expressed VTRNAs (Figure 2C, 4C) further highlights this issue. Although certain tRNA pseudogenes display both high conservation and expression levels, some functional tRNA genes exhibit low conservation and expression (Supplementary Table 1, 2). This observation suggests that approaches like structure-based covariance models and sequence-based hidden Markov models may be more effective in distinguishing functional tRNAs from tRNA pseudogenes (11).

## Conclusion

Accurate classification and naming are crucial, as all human genome studies should adhere to HGNC-approved nomenclature to ensure consistency, clarity, and effective communication across research. This study proposes that certain ncRNA pseudogenes (and functional ncRNAs), such as RNU2-2P, RNU1-27P, and RNU1-28P, may have been misclassified. The combination of high evolutionary conservation and significant expression levels in these pseudogenes suggests they may possess functional potential (59,60).

Although the pseudogenes identified in this study are likely to have a function, computational biology alone cannot confirm this without clinical data and experimental validation. Therefore, future research on ncRNA pseudogenes should focus on analysing clinical data to identify sequence variants associated with diseases and conducting cell line experiments, such as gene knockouts and knockdowns, to establish the functional roles of these pseudogenes.

As our understanding of the significance and functions of ncRNA continues to grow, we anticipate advancements in the quality and availability of ncRNA data, alongside more comprehensive analyses of ncRNA regions to enhance disease diagnosis and support clinical research.

## Supplementary Data

**Supplementary Table 1:** Complete list of PhyloP100 Vertebrate conservation outliers. Black coloured pseudogenes have a z-score > 2. Blue coloured functional ncRNAs have a z-score of < 0.

| Gene Groups | ncRNA outliers for PhyloP100 (Conservation) |
| --- | --- |
| RNU1 | RNU1-28P |
| RNU2 | RNU2-2P, RNU2-4P |
| RNU4 | RNU4-63P |
| RNU5 | RNU5B-5P, RNU5F-1 |
| RNU6 | RNU6-1334P, RNU6-1194P, RNU6-1059P, RNU6-202P, RNU6-693P, RNU6-751P |
| RNU4ATAC | RNU4ATAC15P |
| RNU6ATAC | - |
| RNU11 | - |
| RNU12 | - |
| VTRNA | VTRNA1-1 |
| RNY | - |
| TRNA | TRL-TAA5-1, TRG-CCC7-1, TRG-TCC2-3, TRG-TCC2-4, TRD-GTC2-4, TRD-GTC2-3, TRP-AGG5-1, TRE-TTC15-1, TRK-TTT8-1, TRA-AGC15-1, TRK-TTT14-1, TRP-GGG1-1, TRV-CAC1-3, TRD-GTC2-5, TRSUP-CTA2-1, TRE-CTC1-3, TRG-TCC2-2, TRQ-TTG9-1, TRL-CAA7-1, TRE-CTC1-4, TRV-CAC1-2, TRSUP-CTA3-1, TRD-GTC2-2, TRG-TCC2-5, TRUND-NNN5-1, TRQ-TTG8-1, TRR-CCT6-2, TRE-CTC14-1, TRL-CAG1-5, TRA-TGC10-1, TRL-CAG1-1, TRL-CAG1-4, TRQ-CTG1-3, TRL-AAG1-2, TRV-CAC13-1, TRE-CTC1-5, TRE-CTC1-2, TRS-AGA7-1, TRE-TTC14-1, TRC-GCA7-1, TRL-AAG1-1, TRE-CTC13-1, TRL-CAG1-2, TRD-GTC2-1, TRUND-NNN1-1, TRK-CTT9-1, TRE-CTC15-1, TRE-CTC12-1, TRE-CTC4-1, TRU-TCA2-1, TRD-GTC10-1, TRK-CTT6-1, TRE-CTC11-1, TRE-CTC9-1, TRE-TTC4-2, TRG-CCC4-1, TRN-GTT2-7, TRV-AAC1-3, TRN-GTT2-8, TRV-CAC8-1, TRG-GCC1-4, TRG-TCC4-1, TRL-CAG1-3, TRK-TTT2-1, TRA-AGC9-1, TRL-CAA5-1, TRK-CTT7-1, TRY-ATA1-1, TRK-CTT5-1, TRR-CCT8-1 |
| RN7SL | RN7SL471P |
| RNU7 | RNU7-1 |
| RN7SK | RN7SKP70, RN7SKP253, RN7SKP12, RN7SKP123 |

**Supplementary Table 2:** Complete list of ENCODE gene expression outliers. Black coloured pseudogenes have a z-score > 2. Blue coloured functional ncRNAs have a z-score of < 0.

| Gene Groups | ncRNA outliers for ENCODE (gene expression) |
| --- | --- |
| RNU1 | RNU1-27P, RNU1-28P, RNU1-125P, RNU1-67P, RNU1-72P, RNU1-39P, RNU1-16P, RNU1-34P, RNU1-70P, RNU1-108P, RNU1-91P, RNU1-83P, RNU1-134P, RNU1-109P, RNU1-73P, RNU1-29P, RNU1-103P, RNU1-100P, RNU1-143P, RNU1-68P, RNU1-129P, RNU1-124P, RNU1-89P, RNU1-18P, RNU1-51P, RNU1-93P, RNU1-104P, RNU1-30P, RNU1-19P, RNU1-31P, RNU1-94P, RNU1-5P, RNU1-6P |
| RNU2 | RNU2-2P |
| RNU4 | RNU4-46P, RNU4-9P, RNU4-38P, RNU4-32P, RNU4-78P, RNU4-39P, RNU4-25P, RNU4-59P |
| RNU5 | RNU5E-6P, RNU5B-2P, RNU5D-2P, RNU5B-3P |
| RNU6 | RNU6-1189P, RNU6-82P, RNU6-107P, RNU6-1016P, RNU6-60P, RNU6-853P, RNU6-312P, RNU6-146P, RNU6-485P, RNU6-101P, RNU6-1099P, RNU6-228P, RNU6-396P, RNU6-112P, RNU6-979P, RNU6-118P, RNU6-1088P, RNU6-26P, RNU6-850P, RNU6-230P, RNU6-876P, RNU6-658P, RNU6-1275P, RNU6-1062P, RNU6-1272P, RNU6-275P, RNU6-549P, RNU6-914P, RNU6-467P, RNU6-379P, RNU6-611P, RNU6-638P, RNU6-171P, RNU6-418P, RNU6-1311P, RNU6-1255P, RNU6-174P, RNU6-90P, RNU6-750P, RNU6-1245P, RNU6-33P, RNU6-1053P, RNU6-807P, RNU6-1010P, RNU6-516P, RNU6-94P, RNU6-761P, RNU6-140P, RNU6-1048P, RNU6-808P, RNU6-343P, RNU6-1157P, RNU6-346P, RNU6-130P, RNU6-122P, RNU6-828P, RNU6-530P, RNU6-930P, RNU6-619P, RNU6-1098P, RNU6-513P, RNU6-652P, RNU6-100P, RNU6-444P, RNU6-881P, RNU6-298P, RNU6-1223P, RNU6-540P, RNU6-583P, RNU6-302P, RNU6-49P, RNU6-859P, RNU6-548P, RNU6-1306P, RNU6-1256P, RNU6-322P, RNU6-944P, RNU6-1251P, RNU6-890P, RNU6-762P, RNU6-37P, RNU6-882P, RNU6-531P, RNU6-476P, RNU6-3P, RNU6-5P, RNU6-388P, RNU6-757P, RNU6-577P, RNU6-377P, RNU6-97P, RNU6-446P, RNU6-1118P, RNU6-1217P, RNU6-689P, RNU6-785P, RNU6-998P, RNU6-190P, RNU6-126P, RNU6-1079P, RNU6-681P, RNU6-310P, RNU6-182P, RNU6-479P, RNU6-969P, RNU6-1323P, RNU6-151P, RNU6-185P, RNU6-589P, RNU6-621P, RNU6-1024P, RNU6-1095P, RNU6-722P, RNU6-906P, RNU6-564P |
| RNU4ATAC | RNU4ATAC11P, RNU4ATAC17P, RNU4ATAC16P, RNU4ATAC18P |
| RNU6ATAC | RNU6ATAC24P, RNU6ATAC10P |
| RNU11 | - |
| RNU12 | - |
| VTRNA | - |
| RNY | RNY3P2, RNY3P11, RNY1P14, RNY3P8, RNY1P16, RNY1P12, RNY1P9 |

| TRNA | MT-TF, MT-TL1, NMTRQ-TTG12-1, NMTRS-TGA3-1 |
| --- | --- |
| RN7SL | RN7SL471P, RN7SL4P, RN7SL5P, RN7SL128P, RN7SL81P, RN7SL359P,<br>RN7SL657P, RN7SL731P, RN7SL698P, RN7SL674P, RN7SL648P, RN7SL396P,<br>RN7SL431P, RN7SL192P, RN7SL288P, RN7SL368P, RN7SL449P, RN7SL145P,<br>RN7SL558P, RN7SL246P, RN7SL377P, RN7SL473P, RN7SL735P, RN7SL737P,<br>RN7SL838P, RN7SL738P, RN7SL296P, RN7SL220P, RN7SL664P, RN7SL650P,<br>RN7SL752P, RN7SL404P, RN7SL165P, RN7SL689P, RN7SL153P, RN7SL769P,<br>RN7SL517P, RN7SL381P, RN7SL786P, RN7SL444P, RN7SL521P, RN7SL393P,<br>RN7SL801P, RN7SL394P, RN7SL141P, RN7SL798P, RN7SL280P, RN7SL555P,<br>RN7SL535P, RN7SL68P, RN7SL608P, RN7SL630P, RN7SL398P, RN7SL656P,<br>RN7SL57P, RN7SL709P, RN7SL600P, RN7SL834P, RN7SL15P, RN7SL362P,<br>RN7SL751P, RN7SL356P, RN7SL800P, RN7SL417P, RN7SL329P, RN7SL301P,<br>RN7SL300P, RN7SL23P, RN7SL172P, RN7SL411P, RN7SL321P, RN7SL138P,<br>RN7SL75P, RN7SL861P, RN7SL653P, RN7SL465P, RN7SL242P, RN7SL364P,<br>RN7SL660P, RN7SL728P, RN7SL720P, RN7SL559P, RN7SL809P, RN7SL481P,<br>RN7SL749P, RN7SL105P, RN7SL443P, RN7SL788P, RN7SL118P, RN7SL130P,<br>RN7SL760P, RN7SL836P, RN7SL812P, RN7SL745P, RN7SL308P, RN7SL541P,<br>RN7SL88P, RN7SL200P, RN7SL16P, RN7SL505P, RN7SL30P, RN7SL37P,<br>RN7SL36P, RN7SL566P, RN7SL166P, RN7SL464P, RN7SL173P, RN7SL552P,<br>RN7SL230P, RN7SL146P, RN7SL683P, RN7SL174P, RN7SL127P, RN7SL326P,<br>RN7SL273P, RN7SL334P, RN7SL181P, RN7SL328P, RN7SL452P, RN7SL390P,<br>RN7SL239P, RN7SL513P, RN7SL428P, RN7SL441P, RN7SL655P, RN7SL482P,<br>RN7SL67P, RN7SL851P, RN7SL563P, RN7SL663P, RN7SL344P, RN7SL833P,<br>RN7SL382P, RN7SL124P, RN7SL502P, RN7SL688P, RN7SL19P, RN7SL715P,<br>RN7SL638P, RN7SL767P, RN7SL333P, RN7SL507P, RN7SL587P, RN7SL107P,<br>RN7SL113P, RN7SL237P, RN7SL827P, RN7SL25P, RN7SL569P, RN7SL375P,<br>RN7SL116P, RN7SL8P, RN7SL277P, RN7SL646P, RN7SL672P, RN7SL743P,<br>RN7SL297P, RN7SL775P, RN7SL434P, RN7SL47P, RN7SL177P, RN7SL284P |
| RNU7 | RNU7-40P, RNU7-96P, RNU7-57P, RNU7-84P, RNU7-3P, RNU7-154P, RNU7-124P,<br>RNU7-20P, RNU7-140P, RNU7-151P, RNU7-75P, RNU7-161P, RNU7-149P,<br>RNU7-77P |
| RN7SK | RN7SKP255, RN7SKP95, RN7SKP92, RN7SKP54, RN7SKP16, RN7SKP22,<br>RN7SKP9, RN7SKP26, RN7SKP69, RN7SKP51, RN7SKP239, RN7SKP116,<br>RN7SKP103, RN7SKP154, RN7SKP230, RN7SKP78, RN7SKP110, RN7SKP70,<br>RN7SKP80, RN7SKP195, RN7SKP283, RN7SKP269, RN7SKP11, RN7SKP71,<br>RN7SKP106, RN7SKP151, RN7SKP137, RN7SKP97, RN7SKP56, RN7SKP124,<br>RN7SKP253, RN7SKP176, RN7SKP227, RN7SKP115, RN7SKP296, RN7SKP134,<br>RN7SKP173, RN7SKP292, RN7SKP38, RN7SKP234, RN7SKP36, RN7SKP190,<br>RN7SKP184, RN7SKP288, RN7SKP74, RN7SKP50, RN7SKP43, RN7SKP96,<br>RN7SKP245, RN7SKP248, RN7SKP187, RN7SKP30, RN7SKP102, RN7SKP175,<br>RN7SKP258, RN7SKP25, RN7SKP198, RN7SKP160, RN7SKP127, RN7SKP118,<br>RN7SKP139, RN7SKP185, RN7SKP228, RN7SKP180, RN7SKP221, RN7SKP55,<br>RN7SKP186, RN7SKP273, RN7SKP271 |

**Supplementary Table 3:** Current ncRNA and pseudogene names and suggested reclassification, with random forest functional probability scores.

| HGNC Gene Symbol | Current Classification | Suggested Classification | Random Forest Predicted Functional Probability |
| --- | --- | --- | --- |
| RNU2-2P | Pseudogene | Functional | 1.00 |
| RNU1-27P | Pseudogene | Functional | 1.00 |
| RNU1-28P | Pseudogene | Functional | 1.00 |
| RNU5B-5P | Pseudogene | Functional | 1.00 |
| RNU6-1194P | Pseudogene | Functional | 0.3 |
| RNU6-1334P | Pseudogene | Functional | 0.75 |
| RN7SL471P | Pseudogene | Functional | 0.92 |
| RN7SKP70 | Pseudogene | Functional | 0.58 |
| TRL-TAA5-1 | Pseudogene | Functional | 1.00 |
| RNU6ATAC10P | Pseudogene | Functional | 0.78 |
| MT-TL1 | Functional | Pseudogene | 0.03 |
| NMTRS-TGA3-1 | Functional | Pseudogene | 0.03 |
| TRA-AGC15-1 | Functional | Pseudogene | 0.00 |
| TRA-AGC9-1 | Functional | Pseudogene* | 0.01 |
| TRA-TGC10-1 | Functional | Pseudogene* | 0.00 |
| TRC-GCA7-1 | Functional | Pseudogene* | 0.00 |
| TRD-GTC10-1 | Functional | Pseudogene* | 0.00 |
| TRD-GTC2-1 | Functional | Pseudogene* | 0.00 |
| TRD-GTC2-2 | Functional | Pseudogene* | 0.00 |
| TRD-GTC2-3 | Functional | Pseudogene* | 0.00 |
| TRD-GTC2-4 | Functional | Pseudogene* | 0.00 |
| TRD-GTC2-5 | Functional | Pseudogene* | 0.00 |
| TRE-CTC1-2 | Functional | Pseudogene* | 0.00 |
| TRE-CTC1-3 | Functional | Pseudogene* | 0.00 |
| TRE-CTC1-4 | Functional | Pseudogene* | 0.00 |
| TRE-CTC1-5 | Functional | Pseudogene* | 0.00 |
| TRE-CTC11-1 | Functional | Pseudogene* | 0.00 |
| TRE-CTC12-1 | Functional | Pseudogene* | 0.01 |
| TRE-CTC13-1 | Functional | Pseudogene* | 0.00 |
| TRE-CTC14-1 | Functional | Pseudogene* | 0.00 |
| TRE-CTC15-1 | Functional | Pseudogene* | 0.00 |
| TRE-CTC4-1 | Functional | Pseudogene* | 0.01 |
| TRE-CTC9-1 | Functional | Pseudogene* | 0.00 |
| TRE-TTC14-1 | Functional | Pseudogene* | 0.00 |
| TRE-TTC15-1 | Functional | Pseudogene* | 0.00 |
| TRE-TTC4-2 | Functional | Pseudogene* | 0.00 |
| TRG-CCC4-1 | Functional | Pseudogene* | 0.00 |
| TRG-CCC7-1 | Functional | Pseudogene* | 0.00 |
| TRG-GCC1-4 | Functional | Pseudogene* | 0.01 |
| TRG-TCC2-2 | Functional | Pseudogene* | 0.00 |
| TRG-TCC2-3 | Functional | Pseudogene* | 0.00 |
| TRG-TCC2-4 | Functional | Pseudogene* | 0.00 |
| TRG-TCC2-5 | Functional | Pseudogene* | 0.00 |
| TRG-TCC4-1 | Functional | Pseudogene* | 0.01 |
| TRK-CTT5-1 | Functional | Pseudogene* | 0.19 |
| TRK-CTT6-1 | Functional | Pseudogene* | 0.00 |
| TRK-CTT7-1 | Functional | Pseudogene* | 0.19 |
| TRK-CTT9-1 | Functional | Pseudogene* | 0.00 |
| TRK-TTT14-1 | Functional | Pseudogene* | 0.00 |
| TRK-TTT2-1 | Functional | Pseudogene* | 0.01 |
| TRK-TTT8-1 | Functional | Pseudogene* | 0.00 |
| TRL-AAG1-1 | Functional | Pseudogene* | 0.00 |
| TRL-AAG1-2 | Functional | Pseudogene* | 0.00 |
| TRL-CAA5-1 | Functional | Pseudogene* | 0.18 |
| TRL-CAA7-1 | Functional | Pseudogene* | 0.00 |
| TRL-CAG1-1 | Functional | Pseudogene* | 0.00 |
| TRL-CAG1-2 | Functional | Pseudogene* | 0.00 |
| TRL-CAG1-3 | Functional | Pseudogene* | 0.01 |
| TRL-CAG1-4 | Functional | Pseudogene* | 0.00 |
| TRL-CAG1-5 | Functional | Pseudogene* | 0.00 |
| TRN-GTT2-7 | Functional | Pseudogene* | 0.00 |
| TRN-GTT2-8 | Functional | Pseudogene* | 0.00 |
| TRP-AGG5-1 | Functional | Pseudogene* | 0.00 |
| TRP-GGG1-1 | Functional | Pseudogene* | 0.00 |
| TRQ-CTG1-3 | Functional | Pseudogene* | 0.00 |
| TRQ-TTG8-1 | Functional | Pseudogene* | 0.00 |
| TRQ-TTG9-1 | Functional | Pseudogene* | 0.00 |
| TRR-CCT6-2 | Functional | Pseudogene* | 0.00 |
| TRR-CCT8-1 | Functional | Pseudogene* | 0.07 |
| TRS-AGA7-1 | Functional | Pseudogene* | 0.00 |
| TRSUP-CTA2-1 | Functional | Pseudogene* | 0.00 |
| TRSUP-CTA3-1 | Functional | Pseudogene* | 0.00 |
| TRU-TCA2-1 | Functional | Pseudogene* | 0.00 |
| TRUND-NNN1-1 | Functional | Pseudogene* | 0.00 |
| TRUND-NNN5-1 | Functional | Pseudogene* | 0.00 |
| TRV-AAC1-3 | Functional | Pseudogene* | 0.00 |
| TRV-CAC1-2 | Functional | Pseudogene* | 0.00 |
| TRV-CAC1-3 | Functional | Pseudogene* | 0.00 |
| TRV-CAC13-1 | Functional | Pseudogene* | 0.00 |
| TRV-CAC8-1 | Functional | Pseudogene* | 0.00 |
| TRY-ATA1-1 | Functional | Pseudogene* | 0.19 |

\*Consult with a curator and evaluate using the tRNAscan-SE models to verify function.

